# Multi-representation DeepInsight: an improvement on tabular data analysis

**DOI:** 10.1101/2023.08.02.551620

**Authors:** Alok Sharma, Yosvany López, Shangru Jia, Artem Lysenko, Keith A. Boroevich, Tatsuhiko Tsunoda

## Abstract

Tabular data analysis is a critical task in various domains, enabling us to uncover valuable insights from structured datasets. While traditional machine learning methods have been employed for feature engineering and dimensionality reduction, they often struggle to capture the intricate relationships and dependencies within real-world datasets. In this paper, we present Multi-representation DeepInsight (abbreviated as MRep-DeepInsight), an innovative extension of the DeepInsight method, specifically designed to enhance the analysis of tabular data. By generating multiple representations of samples using diverse feature extraction techniques, our approach aims to capture a broader range of features and reveal deeper insights. We demonstrate the effectiveness of MRep-DeepInsight on single-cell datasets, Alzheimer’s data, and artificial data, showcasing an improved accuracy over the original DeepInsight approach and machine learning methods like random forest and L2-regularized logistic regression. Our results highlight the value of incorporating multiple representations for robust and accurate tabular data analysis. By embracing the power of diverse representations, MRep-DeepInsight offers a promising avenue for advancing decision-making and scientific discovery across a wide range of fields.

## Introduction

Tabular data analysis plays a crucial role in various fields, ranging from biomedical research to finance and beyond. The ability to extract meaningful insights from structured datasets is vital for making informed decisions and driving advancements in diverse domains. Traditional machine learning (ML) methods, such as feature engineering and dimensionality reduction, have long been employed to uncover patterns and relationships within tabular data^1–4^. However, these approaches often struggle to capture the complex interactions and dependencies inherent in real-world datasets^5–7^.

The fundamental approach for ML methods involves processing a column vector of size *d* × 1 to extract pertinent information essential for classification or regression. However, as the complexity of tabular data continues to grow, ML techniques face challenges in identifying relevant class types. Such as accurately determining phenotypes through the processes of feature extraction and classification plays a pivotal role in diagnosis and disease analysis. It is important to note that feature selection is of vital significance across various research domains, extending beyond genomic data analysis. Hence, the procedures of feature selection, feature extraction, and classification significantly impact the reliability of ML algorithms. However, traditional ML approaches disregard crucial neighborhood information and presumptuously assume that each component of a sample is independent of the others.

On the contrary, two-dimensional convolutional neural networks (CNNs) are part of deep learning architectures that have shown immense potential and garnered significant interest in the field of image analysis^3^. What sets CNNs apart from traditional ML methods is their unique approach to feature extraction and classification, accomplished through their convolutional layers that process an input image structured as a *p* × *q* feature matrix. The utilization of CNNs brings forth a plethora of advantages, including efficient feature extraction from spatially coherent pixels, the ability to facilitate deeper networks with fewer parameters through weight sharing^8^, detection of higher-order statistics and nonlinear correlations, and the ability to achieve remarkable performance even with a reduced number of samples.

A key strength of CNNs lies in their adeptness at harnessing the spatial relationships intrinsic to images. By effectively extracting features from adjacent pixels through their layers, CNNs capitalize on the wealth of information shared by nearby pixels. This enables them to capture intricate spatial patterns that traditional ML approaches often overlook. As a result, CNNs present a highly effective solution for tasks where spatial context plays a crucial role in the accurate analysis and classification of data.

With the advent of techniques like DeepInsight^5^, the repertoire of deep learning techniques have expanded for tabular data analysis, offering the promise of automated feature learning and improved predictive accuracy^5,6,9–16,16,17^. DeepInsight has consistently demonstrated remarkable success in uncovering hidden patterns for various kinds of data^9,10,14^ and becoming a part of the winning model in the Kaggle.com competition hosted by MIT and Harvard University^18^. Nevertheless, there remains room for enhancement, as DeepInsight primarily operates on a single representation of the data.

To enhance the capabilities of the DeepInsight suite, we leverage the power of CNNs pretrained with large datasets of images and utilize transfer learning. Transfer learning allows us to benefit from the knowledge learned by CNNs from extensive image datasets, such as ImageNet, without the need to train the entire network from scratch. By initializing the CNNs with these pretrained weights, the model gains the ability to extract high-level and spatial features effectively, even when applied to tabular data represented as images. This integration of transfer learning significantly enhances the feature extraction process within the DeepInsight framework, enabling more robust and accurate analysis of complex tabular datasets.

In this paper, we propose an innovative extension of DeepInsight, termed Multi-representation DeepInsight or MRep-DeepInsight, which aims to overcome the limitations of the original approach by generating multiple representations of samples. By harnessing the power of diverse data representations, we seek to capture a broader range of features and reveal deeper insights within tabular datasets.

The primary motivation behind MRep-DeepInsight is to leverage the complementary nature of different data representations. By simultaneously employing various feature mapping techniques, such as Uniform Manifold Approximation and Projection (UMAP)^19^, t-distributed stochastic neighbor embedding (SNE)^20^, blurring technique^21^, and Gabor filtering^22^, we generate multiple representations of each sample, thereby providing a comprehensive view of its underlying characteristics. These diverse representations are then integrated within the DeepInsight framework, enabling more robust and accurate analysis of tabular data. An overview depicting the MRep-DeepInsight method is shown in Fig. 1 (see Methods for details). The tabular data is converted to various representations in the transformation phase (Fig. 1a). The obtained images are processed via a CNN for training (Fig. 1b). Finally a novel test sample (*t* in Fig. 1c) is given to the trained model for classification into defined categories.

**Figure 1:**
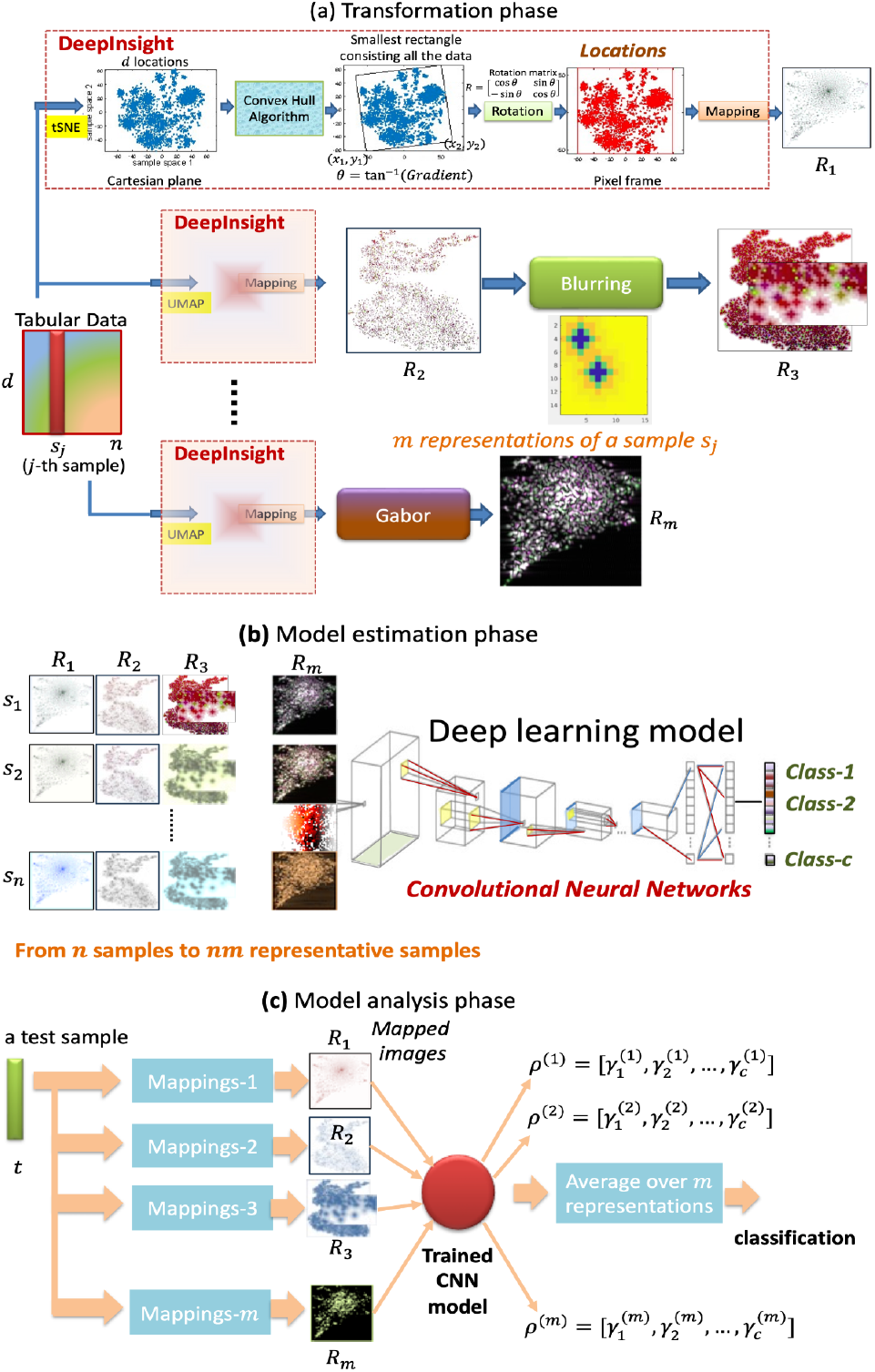
An outline of the MRep-DeepInsight approach. 1a) In the transformation phase, tabular data is converted to image samples using a multi-representation strategy. 1b) Multi-representation samples are processed by a CNN for training. 1c) A novel test sample is analyzed to one of the defined classes.

The MRep-DeepInsight technique can be readily adapted to tabular datasets with improved accuracy. In this paper, we have shown the utility of MRep-DeepInsight on single-cell datasets, Alzheimer’s data, and artificial datasets. By incorporating multiple representations, we aim to capture intricate relationships, interactions, and nonlinearities often overlooked by traditional ML methods or even the original DeepInsight approach.

In this paper, we present a thorough evaluation of MRep-DeepInsight on a diverse set of tabular datasets. The presented results demonstrate the effectiveness of the MRep-DeepInsight approach in improving classification accuracy by uncovering previously hidden patterns. We also discuss the implications of MRep-DeepInsight for various domains that rely on accurate and comprehensive tabular data analysis.

Overall, MRep-DeepInsight represents a significant advancement in the field of tabular data analysis, offering a novel and effective approach for extracting insights from structured datasets. By embracing the power of multiple representations, this approach opens up new possibilities for improved decision-making, scientific discovery, and advancements across a wide range of fields.

## Results

### Experimental setup

To evaluate the effectiveness of the MRep-DeepInsight method, we conducted experiments using six diverse datasets and compared the results with state-of-the-art classifiers. The dataset collection consists of the following: 1 single-cell dataset containing scRNA-seq and ATAC-seq data, 1 single-cell dataset with scRNA-seq only, 1 single-cell dataset with ATAC-seq only, 1 miRNA expression dataset of Alzheimer disease, and 2 artificial datasets (Ringnorm and Madelon). Our primary objective is to demonstrate improved classification accuracy by converting non-image data into multiple representations and processing them using the CNN architecture implemented in the MRep-DeepInsight method. In this study, we utilized two nets, ResNet-50^23^ and EfficientNet-B6^24^.

The datasets were divided into training, validation, and test sets in an 80:10:10 proportion, respectively. The model was trained on the training set, and its fitness was evaluated using the validation set. Hyperparameters were selected to minimize the validation error, particularly for the ResNet-50 architecture. Default hyperparameters were mostly used for the EfficientNet-B6 architecture. Importantly, the test set was completely separated from the training and model fitting stages to ensure an unbiased assessment of the final model’s performance. Classification accuracy, defined as the percentage of correctly classified samples from the test set, was computed as the evaluation metric.

The description of the datasets used in our experiments is as follows: The single-cell data is sourced from PBMC reference, containing scRNA-seq or gene expression profiles with 9602 samples, where scRNAseq consists of 378 genes or dimensions. Additionally, the dataset contains ATAC-seq profiles with corresponding samples and 578 dimensions. This dataset has been obtained from the 10x Chromium Next GEM Single Cell Multiome ATAC + Gene Expression sequencing platform (https://support.10xgenomics.com/single-cell-multiome-atac-gex/datasets/1.0.0/pbmc_granulocyte_sorted_10k) and analyzed using the Cell Ranger ARC pipeline, which includes counting UMIs based on the fastq data (https://support.10xgenomics.com/single-cell-multiome-atac-gex/software/pipelines/latest/what-is-cell-ranger-arc). Notably, batch normalization was not performed on this dataset to preserve the original features. It constitutes 17 distinct cell types, posing a multi-class classification challenge. Among the majority of cell types are CD14 monocyte, CD4 naive, CD8 naive and CD4 TCM (see Supplement Table S4 for the full list of cell types). Notably, CD14 monocyte stands as the predominant cell type in the dataset, accounting for over 26.5% of the samples (same as previously reported^25^). Furthermore, we utilized the Alzheimer’s disease (AD) dataset from a previous study^26^, which comprises 1309 samples of AD and normal controls (NC). Further details of AD dataset can be seen in Supplement Table S5. Lastly, we incorporated two artificial datasets. The first one is Madelon^27^, which consists of 2600 samples and 500 dimensions. It represents a two-class classification problem with continuous input variables, and it exhibits multivariate and highly non-linear characteristics. The second artificial dataset is ringnorm^28^, which is a 20-dimensional, two-class classification problem with 7400 samples. Each class is drawn from a multivariate normal distribution, where class 1 has a zero mean and four times the identity covariance, while class 2 has a mean of a = 2/sqrt(20) with unit covariance. A summary of these datasets can be found in Table 1.

**Table 1.**
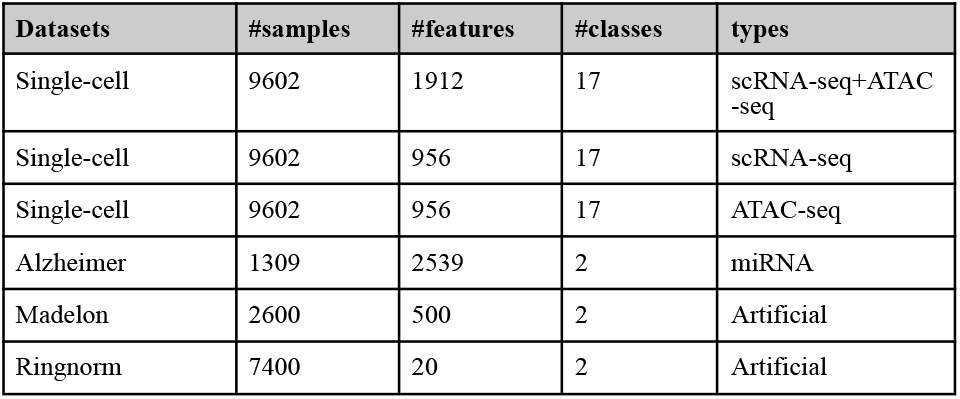
Summary of datasets used.

### Performance comparison

In this section, we compare the performance of MRep-DeepInsight with existing state-of-the-art classifiers, including random forest, L2-regularized logistic regression, (see Section 7 of Supplement File 1 for more details about these techniques) and DeepInsight. To ensure a fair comparison, we optimized the hyperparameters of each method using grid search optimization, except for DeepInsight, which utilized Bayesian optimization. The best-fit models on the validation set were then evaluated on separate test sets.

The purpose of this comparison is to demonstrate how MRep-DeepInsight enhances the characteristics of the data by integrating multiple mapping schemes, leading to improved classification performance. We evaluated the performance on six diverse datasets, and the classification accuracy results are summarized in Table 2 (refer to Supplement Table S2 and Supplement Table S3 for a brief discussion on hyperparameters).

**Table 2.** Classification accuracy (in percentage) comparison across datasets and models.

| Datasets | Random forest | Logistic regression | DeepInsight | MRep-DeepInsight (ResNet50) | Representations used for MRep-DeepInsight ($m$) |
| --- | --- | --- | --- | --- | --- |
| Single-cell: scRNA-seq+ ATAC-seq | 83.2 | 88.5 | 88.2 | 90.5 | *tsne (hamming) + umap + Blurring*<br>($m = 4$) |
| Single-cell: scRNA-seq | 84.5 | 86.1 | 89.2 | 90.0 | *tsne (hamming) + tsne (Euclidean) + umap + Gabor*<br>($m = 6$) |
| Single-cell: ATAC-seq | 71.4 | 77.1 | 78.0 | 79.4 | *tsne (hamming) + tsne (Euclidean) + umap + Gabor*<br>($m = 6$) |
| Alzheimer | 70.8 | 54.2 | 78.1 | 86.2 | *tsne (hamming) + Assignment + umap + Gabor*<br>($m = 6$) |
| Madelon | 82.4 | 88.6 | 84.7 | 89.3 | *tsne (hamming) + Blurring + umap + Gabor*<br>($m = 6$) |
| Ringnorm | 95.7 | 73.0 | 96.4 | 96.5 | *tsne (hamming) + Blurring + umap*<br>($m = 4$) |
| *Average* | *81.3* | *77.9* | *85.8* | *88.6* |  |

On the scRNA-seq+ATAC-seq dataset, MRep-DeepInsight achieved a classification accuracy of 90.5% on the test set, surpassing random forest and DeepInsight by over 7% and 1%, respectively. For the scRNA-seq dataset, MRep-DeepInsight exhibited improved performance compared to random forest and L2-regularized logistic regression, and it slightly outperformed the DeepInsight method. In the case of the ATAC-seq only dataset, MRep-DeepInsight outperformed the second-best method by 1.4%.

When applied to the Alzheimer’s disease dataset, MRep-DeepInsight achieved an accuracy of 86.2%, surpassing the DeepInsight method (78%) as well as L2-regularized logistic regression (54.2%) and random forest (70.8%). These results indicate a significant improvement over state-of-the-art methods. MRep-DeepInsight also demonstrated promising results on both artificial datasets.

Across all six datasets, MRep-DeepInsight achieved an average classification accuracy of 88.6%, which is 7% higher than random forest, approximately 11% higher than L2-regularized logistic regression, and 3% higher than the DeepInsight method. These findings underscore the utility and effectiveness of the MRep-DeepInsight approach.

### Uncertainty analysis/Ablation study

This subsection examines the uncertainty in the obtained results by conducting an ablation study. This study involves varying parameters, including changing the CNN architecture from ResNet-50 to EfficientNet-B6. For ResNet-50, the hyperparameters are tuned using the Bayesian optimization technique while for EfficientNet-B6, default hyperparameter values (L2 penalty as weight decay, momentum = 0.9, and learning rate = 0.001) are used. The mini-batch size and epochs are adjusted based on the memory capacity of the GPU used. The results obtained using EfficientNet-B6 are presented in Supplement Figure S2. Additionally, the effects of varying representation schemes are illustrated in Supplement Table S6 (see Section 5 of Supplement File 1 for more discussion). This ablation study provides insights into the impact of different parameters and representation schemes on the performance of the MRep-DeepInsight methodology, thereby helping to assess its robustness and generalizability.

## Discussions

The MRep-DeepInsight methodology presented offers a novel approach to tabular data analysis by leveraging the power of CNNs and multiple image representations. Our findings demonstrate that MRep-DeepInsight outperforms existing state-of-the-art classifiers, including random forest, L2-regularized logistic regression, and the original DeepInsight method, across various datasets.

One of the key strengths of MRep-DeepInsight lies in its ability to generate multiple representations of non-image data, thereby enriching the information available for classification. By converting tabular data into images using manifold techniques and subsequently utilizing CNN architectures for classification, MRep-DeepInsight captures intricate patterns and fine-grained details that are often overlooked by traditional machine learning methods. This capability is particularly crucial in complex domains such as medical research, where accurately identifying phenotypes and diagnosing diseases is of paramount importance.

Our experimental results demonstrate the effectiveness of MRep-DeepInsight on six diverse datasets. Notably, MRep-DeepInsight achieved significant improvements in classification accuracy compared to other ML methods. On the Alzheimer’s disease dataset, MRep-DeepInsight outperformed random forest and the DeepInsight method by around 16% and 8%, respectively. This clearly indicates its ability to leverage the strengths of gene expression data for improved classification. Similar trends were observed on other datasets, where MRep-DeepInsight consistently surpassed or matched the performance of existing methods.

Furthermore, the utilization of well-established CNN architectures, specifically ResNet-50^23^ and EfficientNet-B6^24^, contributed to the success of MRep-DeepInsight. ResNet-50’s ability to handle deep networks and extract intricate features, combined with EfficientNet-B6’s efficiency and capacity to capture fine-grained details, enabled accurate classification of the multiple image representations generated by MRep-DeepInsight. The hierarchical feature extraction capabilities of these CNN architectures played a crucial role in achieving high classification accuracy. It is worth noting that both ResNet-50 and EfficientNet-B6 are pretrained with large image datasets, and transfer learning has been applied to fine-tune the models for tabular data analysis. This integration of transfer learning significantly enhanced the feature extraction process with the DeepInsight framework, leading to improved classification results.

It is worth noting that the performance of MRep-DeepInsight was evaluated using rigorous experimental setups, including training/validation/test set divisions and hyperparameter optimization. The use of separate test sets ensured an unbiased assessment of the final model’s performance. The hyperparameter optimization process, including grid search optimization and Bayesian optimization, allowed us to fine-tune the models and achieve optimal results.

Overall, the results obtained from our experiments highlight the effectiveness and potential of the MRep-DeepInsight methodology for tabular data analysis. By integrating multiple image representations and leveraging CNN architectures, MRep-DeepInsight offers a robust and accurate approach to classification tasks. The improvements in classification accuracy observed across various datasets demonstrate the utility of MRep-DeepInsight in diverse domains, including single-cell analysis, Alzheimer’s disease diagnosis, and artificial datasets.

While this study presents compelling evidence for the superiority of MRep-DeepInsight, further research is warranted to explore its applicability in other domains and investigate its performance on larger and more diverse datasets. Additionally, future work could focus on optimizing the methodology for efficiency and scalability, as well as exploring potential extensions or adaptations of the approach for other types of data.

In summary, MRep-DeepInsight represents a significant advancement in tabular data analysis by harnessing the power of multiple image representations and CNN architectures. This methodology has the potential to revolutionize the field, enabling accurate classification and providing valuable insights into complex datasets.

## Conclusion

In conclusion, the Multi-representation DeepInsight (MRep-DeepInsight) methodology presented in this study offers a novel and effective approach to tabular data analysis. By generating multiple representations of non-image data and utilizing CNNs, MRep-DeepInsight achieves significant improvements in classification accuracy and provides valuable insights into the underlying data structure. The integration of manifold techniques and the use of well-established pre-trained CNN architectures, namely ResNet-50 and EfficientNet-B6, contribute to the enhanced performance of MRep-DeepInsight.

The experimental results across various datasets demonstrate the superiority of MRep-DeepInsight over state-of-the-art classifiers. The methodology’s ability to convert tabular data into images and leverage CNNs for classification enables the accurate identification of phenotypes and improves disease diagnosis in complex domains. The findings have important implications for single-cell analysis, medical research, and artificial datasets. While MRep-DeepInsight shows promising results, there is still room for further exploration and optimization. Future research can focus on scalability, adapting the methodology to different data types, and comparing it with other advanced techniques to unlock its full potential in real-world applications.

Overall, the MRep-DeepInsight methodology provides a powerful solution for tabular data analysis, enhancing classification accuracy and offering valuable insights into complex datasets. Its versatility and performance improvements make it a valuable tool for researchers seeking to uncover hidden patterns and gain deeper insights from their data, driving advancements in various research domains.

## Methods

This section introduces the MRep-DeepInsight methodology proposed in this study. The model comprises three key components: 1) the transformation of images using multiple DeepInsight representations, 2) the utilization of ResNet-50 and EfficientNet-B6 models as the CNN architecture, and 3) the classification of test samples through average weighting (see Fig. 1). The following subsections outline the step-by-step procedures involved in the MRep-DeepInsight methodology.

### Transformation phase of MRep-DeepInsight

This phase allows the conversion of tabular data to multiple representations of image samples for CNNs in an unsupervised manner. Let us consider a sample represented by **x**_*j*_, where **x** has *d* elements or features (rows), and *j* denotes the samples (columns); i.e., **x** = [*x*_1*j*_,, *x*_2*j*_,…, *x*_*dj*_]^*T*^ Therefore, the training data can be represented as *M* = {**x**_*j*_} for *j* = 1, 2,…, *n*, where *n* is the number of samples. Consequently, the tabular data during the training phase is denoted as *M* ∈ ℜ^*d*×*n*^. The DeepInsight model^5^ is utilized to convert the non-image data *M* into image data *E* = {*e*_*j*_} for *i* = 1, 2,…, *n*. Each image sample, *e*_*j*_ has a size of *p* × *q*. The DeepInsight transformation incorporates manifold techniques such as t-SNE^20^ using different distances (e.g., cosine, Euclidean, Mahalanobis and Chebychev), UMAP^19^, Kernel PCA^29^ and PCA^30^ (see Section 6 of Supplement File 1 for more details about these techniques). These manifold mappings can be further enhanced by applying blurring techniques^21^, Gabor filtering^22^ and the assignment distribution algorithm^31^. The DeepInsight pipeline incorporates several additional steps to enhance the transformation process. These steps include the implementation of the convex hull algorithm, rotation of Cartesian coordinates, determination of pixel locations, and mapping of elements to their respective pixel locations (more discussions on feature mapping are given in Section 3 of Supplement File 1).

When using t-SNE, the algorithm constructs a probability distribution that captures the similarity between pairs of samples. Samples with a higher degree of similarity are assigned higher probabilities, while dissimilar samples receive lower probabilities. This probability distribution is then projected onto a two-dimensional plane, allowing for visualization and analysis. To align the distributions, the Kullback-Leibler divergence is minimized, ensuring that the transformed representations accurately represent the relationships and structure within the data.

It is important to note that these manifold techniques are not only employed for sample visualization, but also for visualization of the genes or elements. To achieve this, the transpose of data *M* is used to determine the pixel locations, *P*. Many of these techniques can project data onto a 2D plane. Thus, if *d* > 2 and *n* > 2, it becomes possible to obtain a 2D framework for *M*. The pixel locations can be obtained using the equation:

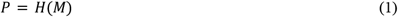

Here, *H* denotes the DeepInsight transform for finding pixel locations, and *M* corresponds to a layer of the training set (e.g. gene expression data). Note that the transpose operation in Eq (1) is not explicitly shown. Once the framework of the pixel locations is determined using Eq (1), the elements can be mapped to generate the corresponding images:

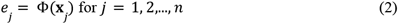

Each non-image sample, **x** ∈ ℜ^*d*^, is mapped to an image sample, *e*_*j*_ ∈ *F*^*p*×*q*^, where *F* is a pixel-coordinates system, Φ is a transformation function (Φ: **x** → *e*), whereas *p* and *q* denote the number of rows and columns, respectively. The transformation, Φ, provides location information for *i*-th element, *x*_*ij*_ ∀*j* = 1, 2… *n*. If the *i*-th location is depicted by [*a*_*i*_, *b*_*i*_], where, *a*_*i*_ is the *i*-th row and *b*_*i*_ is the *i*-th column of the pixel-coordinates, *F*, then *x*_*ij*_ will be mapped at this location, [*a*_*i*_, *b*_*i*_], with its corresponding value.

Different manifold schemes will provide a different Φ transformation. If Φ_*r*_ denotes the *r*-th representative mapping, then Φ_*r*_ : **x** → *e*^(*r*)^ for *r* = 1, 2,…, *m*, i.e, *m* mappings for a sample will be obtained. Mapping all the elements (*i* = 1, 2,…, *d*) corresponding to *j*-th sample, **x**_*j*_, would produce an image sample, 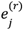. Therefore, we can generalize Eq (2) as:

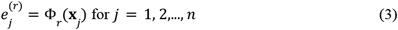

This will, in turn, create *r* locations for the *i*-th element; i.e., Φ_*r*_ contains 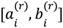 location information for *i* = 1, 2,…, *d*.

The transformation process also includes normalization of the values, typically between the range of [0,1] or [0,255]. In this work, we employed the norm-2 normalization, which was introduced in DeepInsight^5^.

Based on Eq (2), the layer of image data obtained from the transformation is represented as:

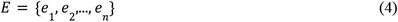

Eq (4) produces one representation (*r* = 1) specific to a manifold method used. For multiple representations (*r* > 1), Eq (4) can be rewritten as:

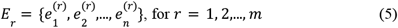

where, *E* ∈ *F*^*p×q×n*^. Here, we obtain *m* representations of the image sample, 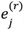, corresponding to a given sample, **x**_*j*_. Hence, by creating *m* representations for each sample, the total number of image samples becomes *nm*.

In this study, a total of *m* = 70 representations can be generated. For more details, see Supplement Figure S1, Supplement Table S1 and Section 1 of Supplement File 1 titled “Multiple representations to find the characteristics of a sample”.

### Model estimation for classification

The classification task in the MRep-DeepInsight methodology is accomplished through the utilization of CNN architectures. Specifically, two well-established CNN models, namely ResNet-50 and EfficientNet-B6, are employed to perform accurate and robust classification of the generated image representations.

ResNet-50 is a deep CNN architecture renowned for its ability to effectively address the challenges associated with training deep networks. With its residual connections, ResNet-50 enables the successful training of models consisting of 50 layers, facilitating the extraction of intricate features from the input images. This architecture has demonstrated exceptional performance across various image analysis tasks, making it a suitable choice for the classification component of MRep-DeepInsight. ResNet-50 leverages the power of transfer learning as it is pretrained with large image datasets, enhancing its feature extraction capabilities and contributing to its robustness and effectiveness^23^.

EfficientNet-B6^24^, on the other hand, is part of the EfficientNet family of CNN architectures known for their superior efficiency and performance. These models achieve a remarkable balance between accuracy and computational efficiency by utilizing a compound scaling technique that optimizes the depth, width, and resolution of the network. EfficientNet-B6 specifically offers a powerful capacity to capture fine-grained details and intricate patterns within image representations. Like ResNet-50, EfficientNet-B6 also benefit from transfer learning, having been pretrained with large image datasets, which significantly enhances its feature extraction capabilities and adds to the overall robustness and effectiveness of MRep-DeepInsight.

By leveraging the capabilities of both ResNet-50 and EfficientNet-B6, the MRep-DeepInsight methodology benefits from the rich hierarchical feature extraction abilities of these CNN architectures, enabling accurate classification of the multiple image representations and providing valuable insights into the underlying structure and characteristics of the data.

Both ResNet-50 and EfficientNet-B6 are supervised models, requiring the inclusion of target or class label information along with samples. To define explicitly, let *X* = {**x**_*j*_} for *j* = 1, 2,…, *n* represent the tabular data with *n* training samples with *d* elements (or features). Additionally, let Ω = {ω_*j*_} be the corresponding class labels, where ω = {1, 2,…, *c*} and *c* is the number of classes. The tabular data *X* is converted into images using Eq (5), resulting in *m* sets, **E** = {*E*_1_, *E*_2_,…, *E*_*m*_}. This procedure generates *nm* images for the *n* training samples, which are then inputted into the CNN architecture for training.

If we define a CNN architecture with its various layers as Ψ, then its output can be depicted as

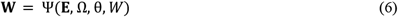

Here, **E** depicts the training set consisting of *nm* images, Ω denotes the class labels of all the samples, θ encompasses the set of all hyperparameters, and *W* represents the weights and biases of the pre-trained CNN model. After training Ψ over several epochs, we obtain updated weights and biases, **W**. This updated model is then utilized for the classification of new samples, enabling accurate predictions based on the learned patterns and representations.

### Model analysis phase

In the model analysis phase, a sample is processed through the trained CNN model (Eq (6)) to obtain predicted probabilities, which are then used to determine its categorization into one of *c* classes. Let us consider an independent test sample, *t*, that was not previously used during the training phase. It is first converted into *m* multiple representations using location information from Eq (3), resulting in *m* images corresponding to *t*_*j*_:

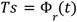

here *Ts* ∈ *F*^*p*×*q*×*m*^ represents the set of *m* mapped images *Ts* = {*R*_1_, *R*_2_,…, *R*_*m*_}, where each image is denoted as *R*_*j*_ ∈ *F*^*p*×*q*^. These *m* images are then fed into the trained CNN model, and each image *R*_*j*_ retrieves a probability distribution from the model:

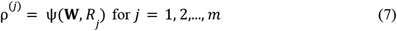

where ψ provides *c* probabilities for the image *R*_*j*_ from the trained CNN model, represented as 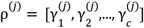. In order to classify the sample, *t*, we can calculate the average probability from Eq (7) to determine the class label:

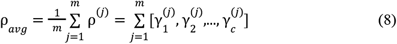

The class label can now be determined from Eq (8) as:

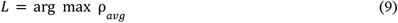

where *L* in Eq (9) depicts the class label assigned to the sample, *t*, based on the highest average probability; i.e., *L* = arg(*t*). This completes the procedure, providing the classification of the test sample using the MRep-DeepInsight methodology.

## Data availability

The single cell data can be downloaded from the link https://support.10xgenomics.com/single-cell-multiome-atac-gex/datasets/1.0.0/pbmc_granulocyte_sorted_10k. The Alzheimer’s disease (AD) dataset is available from the Gene Expression Omnibus (GEO) database at the National Center for Biotechnology Information (NCBI) (https://www.ncbi.nlm.nih.gov/geo/). The exact configuration of AD data is also available at GitHub link https://github.com/alok-ai-lab/MRep-DeepInsight/blob/main/Data/dataset4.mat. Madelon dataset is available from UCI repository https://archive.ics.uci.edu/dataset/171/madelon and Ringnorm dataset is available from the University of Toronto repository https://www.cs.toronto.edu/~delve/data/ringnorm/desc.html.

## Code availability

MRep-DeepInsight is available as a MATLAB package, the latest version can be downloaded from the link https://github.com/alok-ai-lab/MRep-DeepInsight. The ReadMe file with examples is also given to guide a user to execute the codes.

## Acknowledgements

This work was funded by JSPS KAKENHI (JP20H03240), Japan and JST CREST (JPMJCR2231), Japan.

## Author contributions

AS perceived, designed the mapping and classification models, and wrote the first draft and contributed in the subsequent versions of the manuscript. YL designed the EfficientNet-B6 model for MRep-DeepInsight. SJ extracted single-cell datasets. AL improved on the designed models and contributed to the manuscript. KAB aided technical development of the model and helped in the manuscript writeup. TT perceived and contributed to the manuscript writeup. All authors read and approved the manuscript.

