## Supplement File 1 for "Multi-representation DeepInsight: an improvement on tabular data analysis"

### Supplementary File 1

#### 1. Multiple representations to find the characteristics of a sample

The MRep-DeepInsight methodology offers flexibility in utilizing various representations to capture the unique characteristics of samples. In Supplement Figure S1, we present a visualization of 70 different representations that can be employed within the MRep-DeepInsight framework. It is important to note that the methodology is not restricted to these 70 representations alone, and additional relevant representations can be incorporated to further enhance the understanding of sample characteristics. In the current context, we consider the integration of $m = 70$ representations within the MRep-DeepInsight methodology, allowing for a comprehensive analysis of the data.

The MRep-DeepInsight pipeline builds upon the DeepInsight architecture[1] and involves the transformation of tabular data into image samples for CNN application. The pipeline incorporates several key steps, including element arrangement using manifold techniques, application of the convex hull algorithm, rotation in Cartesian coordinates, translation to pixel coordinates, and mapping of original elements to these pixel locations. The manifold technique options available for MRep-DeepInsight include t-SNE[2], UMAP[3], kernel PCA[4], and PCA[5]. In addition, Gabor[6], Blurring scheme[7] and Assignment distribution algorithm[8] will use the representations of manifold techniques to provide modified representations. For instance, if manifold techniques like t-SNE (using hamming distance) and UMAP are selected, then the number of representations, $m$, would be 2. Furthermore, if the Blurring technique is added then it will produce Blurring image for t-SNE (hamming) as well as Blurring image for UMAP. Therefore, total representation for t-SNE (hamming), UMAP and Blurring technique would become $m = 4$. If both the Blurring technique and Assignment are switched on for a manifold technique (such as t-SNE), then it will produce 4 representations, one for t-SNE, one for Blurring image of t-SNE, one is Assignment of t-SNE and the last one is the Blurring image of Assignment image (previously generated). Some more examples of types used in MRep-DeepInsight and the corresponding numbers of representations are given in Supplement Table S1.

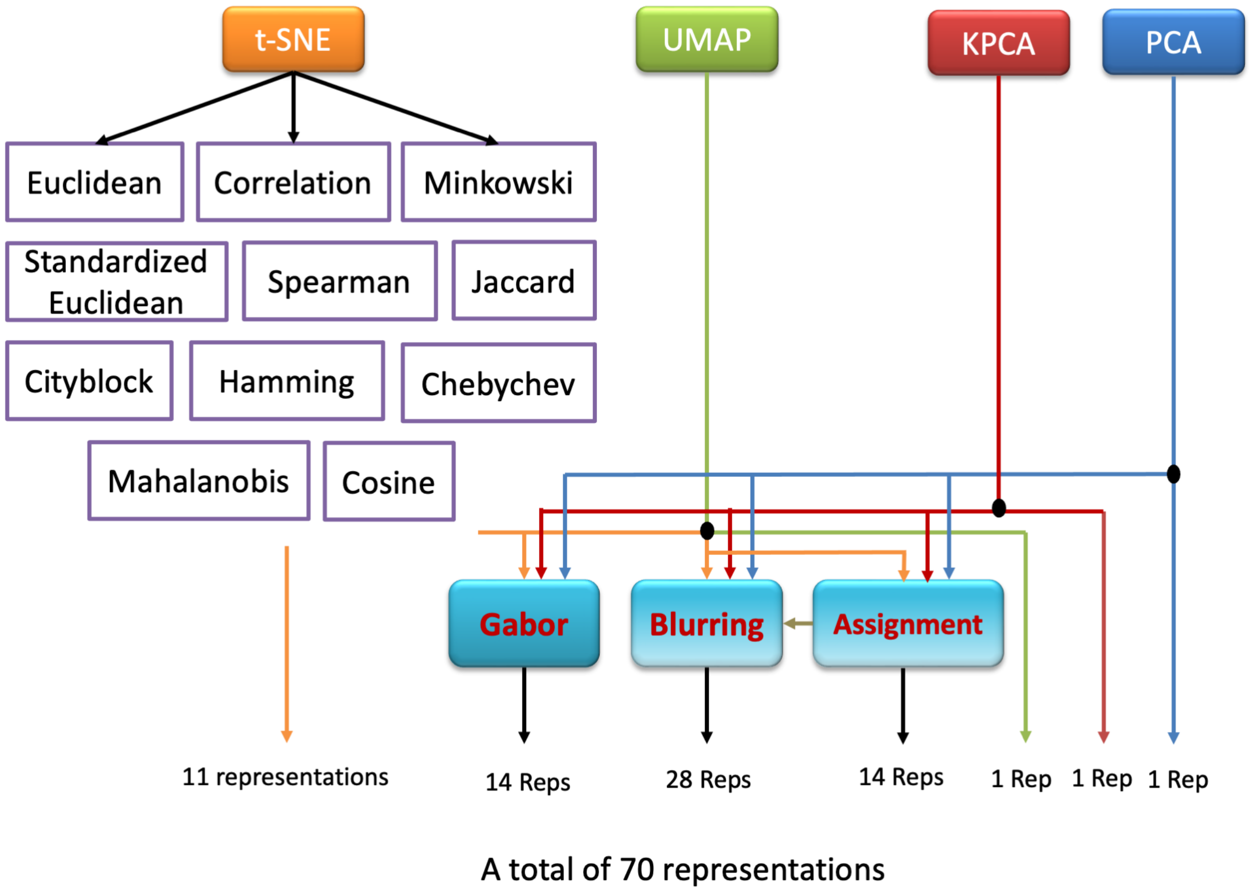


**Supplement Figure S1**: Many representations that can be used by MRep-DeepInsight methodology

**Supplement Table S1**: Examples of Types and the number of representations generated.

| *Type* | *#Representations (m)* |
|---|---|
| t-SNE (hamming) | 1 |
| UMAP | 1 |
| t-SNE (hamming) + UMAP | 2 |
| t-SNE (hamming) + t-SNE (Euclidean) + UMAP | 3 |
| t-SNE (hamming) + UMAP + Blurring | 4 |
| t-SNE (hamming) + UMAP + Blurring + Gabor | 6 |
| t-SNE (hamming) + UMAP + Blurring + Gabor + Assignment | 10 |
| t-SNE (hamming) + Blurring + Assignment | 4 |
| t-SNE (hamming) + t-SNE (Euclidean) + UMAP + Assignment + Blurring + Gabor | 15 |

The expression for the number of representation, $m$, and type can be derived in the following manner. Let $|.|$ be the cardinality function for manifold techniques. Since only t-SNE can have 11 distances (see Supplement Figure 1), we have,

$0 \leq |tSNE| \leq 11$

where, $|tSNE| = 0$, means that t-SNE has not been used. For all other manifold techniques (UMAP, KPCA and PCA) the cardinality would be either 0 (if not used) or 1 (if used). Therefore, it gives,

$|UMAP| = [0, 1]$
$|KPCA| = [0, 1]$
$|PCA| = [0, 1]$

Further, let us define the cardinality for Blurring technique, Assignment and Gabor be $B = |Blurring|$, $A = |Assignment|$ and $G = |Gabor|$. The values of $B$, $A$ and $G$ are either 0 (if not used) or 1 (if used), such as:

$B = [0, 1]$, $A = [0, 1]$, and $G = [0, 1]$

If we define the cardinality of all manifold technique as $|Mt|$, then it can be given as

$$|Mt| = |tSNE| + |UMAP| + |KPCA| + |PCA| \quad \text{(S1)}$$

Eq (S1) computes the number of representations due to the manifold techniques used in MRep-DeepInsight method. Next, the number of representations, $m$ due to $|Mt|$ and Assignment can be given as,

$$m = A \times |Mt| + |Mt| \quad \text{(S2)}$$

If Blurring technique is included then $m$ in Eq (S2) will become

$$m = B(A \times |Mt| + |Mt|) + A \times |Mt| + |Mt| \quad \text{(S3)}$$

From Eq (S3), we can see that Blurring technique will render images for manifold techniques and also for Assignment technique. Furthermore, if Gabor technique is included, the Eq (S3) will become

$$m = B(A \times |Mt| + |Mt|) + A \times |Mt| + |Mt| + G \times |Mt| \quad \text{(S4)}$$

By solving Eq (S4), we get

$$m = (BA + B + A + G + 1)|Mt| \quad \text{(S5)}$$

Considering all possibilities of manifold techniques, the maximum value of $|Mt| = 14$. Moreover, if Gabor, Assignment and Blurring techniques are used then, $m = 5 \times 14 = 70$.

When utilizing t-SNE, various distance measures such as 'Euclidean', 'cosine', 'hamming', and 'correlation', among others, can be employed (as shown in Supplement Figure 1). In cases where the number of elements or data dimensionality exceeds 5000, the Burnes-Hut algorithm is utilized for faster processing, while the exact algorithm is employed for lower-dimensional data.

### 2. CNN parameters

Two CNN implementations used in this study are ResNet-50 and EfficientNet-B6. The details are given below.

*ResNet-50*

The hyperparameters such as learning rate, momentum, and L2 regularization are tuned using the Bayes optimization technique from a range of values. The description of these parameters is given in Supplement Table S2. The hyperparameters are obtained by optimizing the validation error.

**Supplement Table S2**: ResNet-50 training options.

| *Variables* | *Values/range* |
|---|---|
| Net | ResNet-50 |
| Training option | Sgdm<br>(Stochastic gradient descent with momentum) |
| InitialLearningRate | [1e-5 1e-1] |
| Momentum | [0.8 0.95] |
| L2regularization | [1e-10 1e-2] |
| Max Objectives* | 25 |
| MaxTime | 50x60x60 |
| Execution environment | multiple-GPU |
| MaxEpochs* | 50 |
| Min batch size | 1024 |
| Shuffle | Every-epoch |
| Augmentation | Yes |
| Image Size | 224 x 224 |

** First Bayesian optimization technique applied by keeping Maximum Objective to be 25 at 50 Epochs. Once the hyperparameters are found in this way then again, the model is run with obtained hyperparameters for 100-400 Epochs.*

EfficientNet-B6

The hyperparameters for EfficientNet-B6 are depicted in Supplement Table S3.

**Supplement Table S3**: EfficientNet-B6 training options.

| *Variables* | *Values/range* |
|---|---|
| Net | EfficientNet-B6 |
| Training option | SGD (stochastic gradient descent) |
| LearningRate | 0.001 |
| Momentum | 0.9 |
| L2 penalty | 0 |
| Max Objectives | 1 |
| Execution environment | multiple-GPU |
| MaxEpochs | 500 |
| Min batch size | 32 |
| Shuffle | Every-epoch |
| Augmentation | Yes |
| Image Size | 224 x 224 |

Two types of norms are introduced in DeepInsight[1]. However, in this work, we applied Norm-2.

3. **Feature mapping**

In this subsection, a set of pixel locations is obtained from manifold techniques, and all the samples are mapped to these locations. This process is repeated for different manifold techniques, resulting in multiple pixel locations and representations for each sample. Consequently, the tabular data is transformed into multiple representations of a 2D image. The mapping of elements assigns values or shades to these locations. The determination of element locations is based on the training set, and in cases where multiple elements occupy the same spot, their averaged values are mapped onto that location. For example, if locations of elements $g_i$, $g_j$ and $g_k$ are the same $(r, c)$, then the average value $(g_i + g_j + g_k)/3$ is mapped to that location. This process can lead to quantized compression[9], particularly when more than two elements share the same location. The validation and test sets utilized the same pixel locations obtained from the training set.

When converting a tabular sample to an image, an optimum pixel frame size $p \times q$ is obtained. Non-quantized compression occurs when the desired frame size $A \times B$ is greater than $p \times q$ ($p \leq A$ and $q \leq B$). Quantized compression, on the other hand, can be achieved by adjusting the horizontal and vertical sizes of the pixel frame, where $p > A$ and $q > B$. In quantized compression, it is possible for more than two elements share the same location, resulting in the average value of those elements being mapped to that particular location. This phenomenon is referred to as the "overlapping issue". Minimizing the overlapping issue us desirable, especially for that element or feature selection process.

### 4. Dataset configuration

The description of the several datasets used in this work is given below.

*Single-cell data*

The single-cell dataset employed in this study encompasses two profiles: scRNA-seq and ATAC-seq. The scRNA-seq profile has a dimensionality of 378, while the ATAC-seq profile has a dimensionality of 578. It is important to note that the samples and cell types used in both profiles are the same, and further details can be found in Supplement Table S4.

**Supplement Table S4** Single-cell dataset (scRNA-seq and ATAC-seq)

| Cell types | # Total samples |
|---|---|
| CD14 monocyte | 2554 |
| CD4 naive | 1382 |
| CD8 naive | 1354 |
| CD4 TCM | 1113 |
| CD16 monocyte | 442 |
| NK | 403 |
| CD8 TEM_1 | 322 |
| CD8 TEM_2 | 315 |
| Intermediate B | 300 |
| Memory B | 298 |
| CD4 TEM | 286 |
| cDC | 180 |
| Treg | 157 |
| gdT | 143 |
| MAIT | 130 |
| Naïve B | 125 |
| pDC | 98 |

*Alzheimer's Disease data*

The Alzheimer's disease dataset used in this study was sourced from our previous work[10]. The dataset comprises miRNA expression profiles of 1601 Japanese individuals, among which 1021 samples are associated with Alzheimer's disease (AD), while 288 samples represent normal controls (NC). To ensure unbiased evaluation, the samples in the dataset were randomly organized into training and test sets. The dataset contains a total of 2539 miRNA expressions, making it a comprehensive resource for studying Alzheimer's disease.

The test set was constructed by randomly selecting 10% of the samples from the entire dataset. Supplement Table S5 provides detailed information about the Alzheimer's disease dataset, including sample characteristics and other relevant details. By utilizing this valuable dataset, we aim to assess the performance of our MRep-DeepInsight methodology in accurately classifying Alzheimer's disease samples and distinguishing them from normal controls.

**Supplement Table S5**: Alzheimer's disease miRNA dataset

| Phenotype | #total samples | #test samples |
|---|---|---|
| AD | 1021 | 102 |
| NC | 288 | 29 |

### 5. Uncertainty Analysis/Ablation Study

To comprehensively assess the robustness and uncertainty of the MRep-DeepInsight model, we conducted an ablation study by varying its parameters and architecture. This study aims to evaluate how different choices in the Convolutional Neural Network (CNN) architecture and representation types impact the performance of MRep-DeepInsight. The results provide insights into the model's generalization capabilities and its effectiveness in tabular data analysis.

*Ablation Study on CNN Architecture*:

As part of the ablation study, we investigated the impact of changing the CNN architecture from ResNet-50 to EfficientNet-B6. We utilized default hyperparameters of L2 regularization, momentum, and initial learning rate for EfficientNet-B6, while employing Bayesian optimization for tuning hyperparameters in the case of ResNet-50. The miniBatchSize and Epochs were adjusted based on the memory capacity of the GPU used. The results of this comparison are presented in Supplement Figure S2.

In the ablation study, we assessed the classification accuracy achieved by MRep-DeepInsight with EfficientNet-B6 and compared it against the results obtained with ResNet-50 and other competing methods. It can be observed from Supplement Figure S2, that both ResNet-50 with Bayesian optimization technique (BOT) and EfficientNet-B6 without BOT can achieve high performance. This analysis allows us to gain insights into the impact of architecture choice on the overall performance of our method and provides a comprehensive understanding of its generalization capabilities across different parameter settings. The findings from this ablation study contribute to our confidence in the reliability and effectiveness of the MRep-DeepInsight model for tabular data analysis.

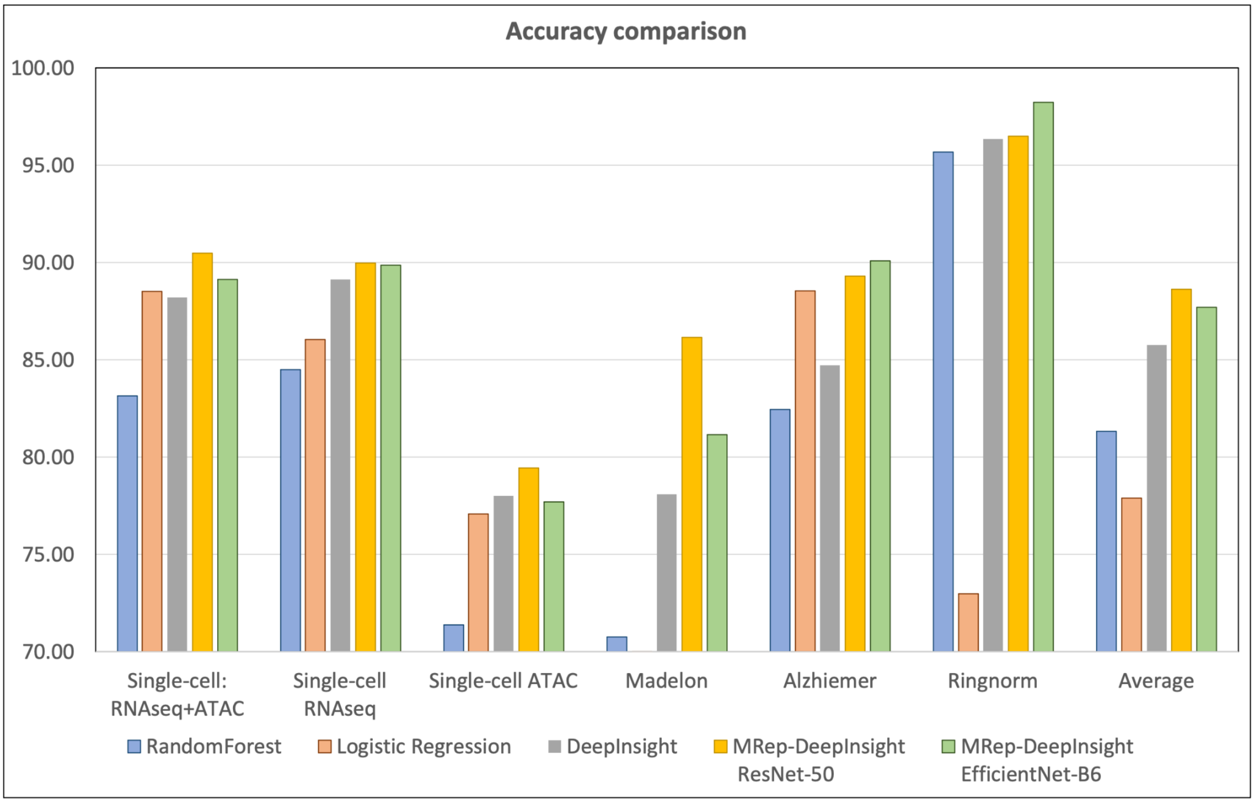


**Supplement Figure S2**: Classification Accuracy of MRep-DeepInsight with EfficientNet-B6 and Comparison with Competing Methods.

*Impact of Representation Types on Performance*

The MRep-DeepInsight method utilizes various types of representations based on different manifold techniques and t-SNE distances. To assess the influence of representation types on performance, we gradually augmented the number of representations (m). The performance of the model was evaluated using the ResNet-50 CNN with hyperparameters tuned in two stages. First, the Bayesian optimization technique was used with a maximum of 25 objectives at 50 Epochs. In the second stage, the best hyperparameters were tuned with 400 Epochs to ensure reduced training time.

Supplement Table S6 depicts the performance of MRep-DeepInsight with different numbers of representations ($m$). We observed that integrating multiple representations, especially with $m = 1$ to 6, increased the classification performance from 84.73% to 89.31%. This highlights the importance of employing multiple representation types to enrich the model's understanding of the data. However, continuously increasing the number of representations (e.g., $m = 15$) had an adverse effect on performance, indicating the significance of finding a moderate value for $m$.

*Recommendation for m value*

Based on the ablation study results, we recommend using a moderate value of $m$ for the MRep-DeepInsight model. A balanced selection of representation types enhances the model's ability to capture the unique characteristics of samples effectively. Striking the right balance between representation diversity and complexity is crucial for achieving optimal classification performance.

In summary, the ablation study on the MRep-DeepInsight model provides valuable insights into its robustness and generalization capabilities. By exploring various CNN architectures and representation types, we gain a deeper understanding of how these choices impact the model's performance. These findings contribute to our confidence in the reliability and effectiveness of the MRep-DeepInsight method for accurate and comprehensive tabular data analysis.

**Supplement Table S6:** Observing the classification performance while changing the types in MRep-DeepInsight method

| Type | #Representations (m) | Classification accuracy |
|---|---|---|
| tsne (hamming) | 1 | 84.73 |
| tsne (hamming) + umap | 2 | 87.02 |
| tsne (hamming) +umap + Assignment | 4 | 87.02 |
| tsne (hamming) + umap + Assignment + Gabor | 6 | 89.31 |
| tsne (hamming) + umap + Assignment + Gabor + Blurring | 10 | 89.31 |
| tsne (hamming) + tsne (Euclidean) + umap + Assignment + Gabor + Blurring | 15 | 85.5 |

### 6. <u>Manifold Techniques</u>

This subsection briefly covers the manifold techniques applied for MRep-DeepInsight methodology. These techniques are t-SNE, Kernel PCA (KPCA), PCA and UMAP. The following sections describe these techniques (except PCA).

#### 6.1. <u>t-SNE</u>

T-distributed stochastic neighbor embedding (t-SNE) is a powerful technique used for visualizing high-dimensional data in a two or three-dimensional space, facilitating the clustering of samples. It achieves this mapping through a non-linear transformation, where similar samples are positioned close to each other while dissimilar samples are placed further apart. Compared to other linear dimensionality reduction methods, t-SNE offers improved visualizations, preserving the topology of the data and enabling a better understanding of complex structures in a lower-dimensional space.

The t-SNE algorithm operates in two main steps. Firstly, it constructs a probability distribution over pairs of samples, assigning higher probabilities to similar sample pairs and lower probabilities to dissimilar pairs. In the second step, it seeks to find a similar probability distribution in a 2D plane and minimizes the Kullback-Leibler divergence between the distributions in the higher-dimensional and lower-dimensional spaces using gradient descent.

In this study, we utilized multiple distances for computing probabilities in t-SNE, although it typically employs the Euclidean distance. The conditional probability, $p_{j|i}$, represents the probability that a sample $x_i$ will select $x_j$ as its neighbor under the Gaussian distribution. Similarly, the conditional probability in the 2D plane, $q_{j|i}$, for mapped samples $y_i$ and $y_j$, is determined based on the Gaussian distribution with a variance. The expressions are defined as:

$$p_{j|i} = \frac{exp(-||x_i - x_j||^2/2\sigma_i^2)}{\sum_{k \neq i} exp(-||x_i - x_k||^2/2\sigma_i^2)}, \text{ and}$$

$$q_{j|i} = \frac{(1+||y_i - y_j||^2)^{-1}}{\sum_{k \neq l} (1+||y_k - y_l||^2)^{-1}}$$

Where $x \in \Re^d$, $y \in \Re^2$, $\sigma_i$ is the variance of the Gaussian that is centered at sample $x_i$, $\sigma_{yi}$ is the variance set to $1/\sqrt{2}$, and $p_{i|i} = 0$.

The optimization process aims to minimize the Kullback-Leibler divergence, ensuring that the mapped samples correctly model the similarity between the higher-dimensional samples. The cost function C is used for this optimization, measuring the difference between the conditional probability distributions in both the higher- and lower-dimensional spaces as

$$C = \sum_i KL(P_i||Q_i) = \sum_i \sum_j log\frac{p_{j|i}}{q_{j|i}}$$

Through t-SNE optimization, mapped samples are positioned in the lower-dimensional space to represent similarities between the samples in the higher-dimensional space. This enables enhanced visualization and clustering, making t-SNE a valuable tool in the MRep-DeepInsight methodology for capturing sample characteristics and uncovering meaningful patterns in the data.

### 6.2 Kernel PCA

Kernel Principal Component Analysis (KPCA) is a powerful extension of the traditional Principal Component Analysis (PCA) technique, widely used for dimensionality reduction and various applications such as visualization, novelty detection, and image de-noising. KPCA incorporates kernel functions to efficiently compute principal components in higher-dimensional spaces without explicitly transforming the data to those spaces.

In KPCA, a projection function $\phi$ is used to map the original data samples $x \in \Re^d$ to a higher-dimensional feature space, which may even be of infinite dimension. However, instead of directly computing the feature space, the kernel trick is applied to obtain mapped samples $y \in \Re^h$, where $h < d$, in a computationally efficient manner.

Assuming the projected dataset with $N$ samples $\phi(x_1), \phi(x_2),..., \phi(x_N)$ is centered (i.e., its mean is zero), the covariance matrix $\Sigma$ can be expressed as $\Sigma = \frac{1}{N}\Phi\Phi^T$ , where $\Phi$ is the data matrix defined as $\Phi = \phi(x_1), \phi(x_2),..., \phi(x_N)$. The eigenvalue decomposition (EVD) of $\Sigma$, denoted as $\Sigma v_i = \lambda_i v_i$ for $i = 1, 2,..., d$, yields eigenvectors $v_i$ and corresponding eigenvalues $\lambda_i$. The eigenvectors $v_i$ can be represented as linear combinations of $\phi(x_1), \phi(x_2),..., \phi(x_N)$, such that $v_i = \Phi u_i$, where $u_i$ is an N-dimensional column vector.

By using the kernel definition $K = \Phi^T\Phi$, the EVD equation ($\Sigma v_i = \lambda_i v_i$) can be expressed as $\frac{1}{N}K u_i = \lambda_i u_i$, where $u_i$ is the eigenvector of the $N \times N$ kernel matrix $K$ corresponding to eigenvalue $\lambda_i$. To ensure normalization, it holds that,

$$1 = v_i^T v_i = u_i^T \Phi^T \Phi u_i = u_i^T K u_i = N\lambda_i u_i^T u_i$$

To perform dimensionality reduction, the projected samples $y_i$ can be obtained from $Y_i = v_i^T \Phi = u_i^T \Phi^T \Phi = u_i^T K$. However, in practice, the projected data does not always have a zero mean. To address this, centralization is applied, and the kernel matrix K is modified to $\hat{K}$ as

$$\hat{K} = K - 1_N K - K 1_N + 1_N K 1_N,$$

Where $1_N$ is an $N \times N$ matrix with all elements set to $1/N$.

### 6.3 UMAP

Uniform Manifold Approximation and Projection (UMAP) is a powerful nonlinear dimensionality reduction technique used for visualizing high-dimensional data in a lower-dimensional space. It is particularly effective in capturing complex data structures and preserving local relationships between data points. UMAP aims to find a faithful representation of the data in a lower-dimensional space while maintaining the global and local structure present in the original high-dimensional data.

The UMAP algorithm builds upon the concept of constructing a fuzzy topological representation of the data, which is then optimized to minimize the difference between the high-dimensional pairwise distances and the pairwise distances in the lower-dimensional space. This optimization process ensures that data points that are close together in the original space remain close in the lower-dimensional embedding.

UMAP uses a combination of fuzzy simplicial sets and Riemannian geometry to achieve its objective. The algorithm starts by creating a fuzzy topological representation of the data, where each data point is connected to its neighbors with fuzzy simplicial sets, allowing for uncertainty in the neighborhood relationships. This fuzzy topological representation is then converted into a low-dimensional Riemannian manifold, where the distances between points are measured based on geodesic distances.

The optimization process involves finding the embedding that minimizes the cross-entropy between the fuzzy simplicial sets in the high-dimensional and lower-dimensional spaces. By minimizing the cross-entropy, UMAP effectively ensures that data points with high similarity are closely mapped in the lower-dimensional embedding.

UMAP claims to offer several advantages over other dimensionality reduction techniques, such as t-SNE and traditional PCA. It is computationally efficient, scalable to large datasets, and exhibits robustness to varying data densities. Moreover, UMAP does not suffer from a "crowding problem," a common issue faced by t-SNE, where data points are disproportionately crowded in the embedding.

Overall, UMAP is a versatile and powerful tool for data visualization and dimensionality reduction, widely used in various domains, including machine learning, bioinformatics, and data analysis. Its ability to preserve local and global structures of the data while efficiently handling large datasets makes it an attractive choice for exploring and understanding complex high-dimensional datasets.

### 7. **Competitive Machine learning techniques**

Here we briefly discuss the two competitive machine learning techniques, random forest and L2-regularized logistic regression.

#### 7.1 L2-regularized logistic regression

Here we developed an L2-regularized logistic regression model to address the classification task. For each of the $c$ classes, an individual model is created with the following formulation:

$$\min_{w} \frac{1}{2}w^T w + C \sum_{j=1}^{d} \log(1 + e^{-y_j w^T x_j})$$

Here, $\frac{1}{2}w^T w$ represents the L2 regularization term (Ridge) which prevents overfitting and improves the generalization of the model, $i = 1, 2, ..., c$ and $d$ is the number of elements or features in the dataset. The logistic regression model estimates the probability of the input sample $x_j$ belonging to class $y_j$. We create models for all the $c$ classes and therefore get coefficients $w_j^i$ (where $j = 1, 2, ..., d$).

For binary classification problems where $c = 2$, only one model is created.

The implementation of L2-regularized logistic regression was achieved using the liblinear package in MATLAB, a powerful tool that offers efficient solutions for large-scale linear classification. For further details, please refer to the liblinear website:
https://www.csie.ntu.edu.tw/~cjlin/liblinear/.

#### 7.2. Random Forest

Random Forest is an ensemble learning method widely used for classification and regression tasks. It operates by constructing multiple decision trees during the training phase. Each tree is trained on a random subset of the data, and during the prediction phase, the ensemble of trees votes to determine the final output. This approach leads to improved accuracy, robustness against overfitting, and enhanced generalization performance.

Random Forest combines the concepts of bagging and feature randomness. Bagging involves bootstrap sampling of the training data, creating diverse subsets for individual tree training. Feature randomness entails randomly selecting a subset of features at each split, ensuring that different trees use different subsets of features.

The final prediction is based on the majority vote of the individual decision trees, making Random Forest a powerful and versatile classification method. It can handle high-dimensional datasets, maintain good performance on large data samples, and effectively deal with missing values and noisy data.

For Random Forest, we employed the TreeBagger package from MATLAB, a tool that efficiently implements the Random Forest algorithm and offers various tuning parameters to optimize the performance of the model.
